# The Effects of Different Vibration Frequencies, Amplitudes and Contraction Levels on Lower Limb Muscles during Graded Isometric Contractions Superimposed on Whole Body Vibration Stimulation

**DOI:** 10.1101/317602

**Authors:** Amit N. Pujari, Richard D. Neilson, Marco Cardinale

## Abstract

**Background:** Indirect vibration stimulation i.e. whole body vibration or upper limb vibration, has been suggested increasingly as an effective exercise intervention for sports and rehabilitation applications. However, there is a lack of evidence regarding the effects of whole body vibration (WBV) stimulation superimposed to graded isometric contractions superimposed on. For this scope, we investigated the effects of WBV superimposed to graded isometric contractions in the lower limbs on muscle activation. We also assessed the agonist-antagonist co-activation during this type of exercise.

Twelve healthy volunteers were exposed to WBV superimposed to graded isometric contractions, at 20, 40, 60, 80 and 100% of the maximum voluntary contractions (V) or just isometric contractions performed on a custom designed horizontal leg press Control (C). Tested stimulation consisted of 30Hzand 50Hz frequencies and 0.5mm and 1.5mm amplitudes. Surface electromyographic activity of Vastus Lateralis (VL), Vastus Medialis (VM) and Biceps Femoris (BF) were measured during V and C conditions. Co-contraction activity of agonist-antagonist muscles was also quantified. The trials were performed in random order.

**Results:** Both the prime mover, (VL) and the antagonist, (BF) displayed significantly higher (P < 0.05) EMG activity with the V than the C condition. For both the VL and BF, the increase in mean EMGrms values depended on the frequency, amplitude and muscle contraction level with 50Hz-0.5mm stimulation inducing the largest neuromuscular activity. 50Hz-0.5mm V condition also led to co-activation ratios significantly (P< 0.05) higher at 40, 80 and 100% of MVC than the C condition.

**Conclusions:** Our results show that the isometric contraction superimposed on vibration stimulation leads to higher neuromuscular activity compared to isometric contraction alone in the lower limbs. Compared to the control condition, the vibratory stimulation leads to higher agonist-antagonist co-activation of the muscles around the knee joint in all vibration conditions and effort levels. The combination of vibration magnitude (frequency and amplitude) and the level of muscle contraction affect neuromuscular activity rather than vibration frequency alone. Results of this study suggest that more parameters need to be taken into consideration when designing vibration exercise programs for sports and rehabilitation purposes.

## Background

Vibration stimulation has been used as a diagnostic tool in neurological studies since the 70s [1]. Vibration has also been studied extensively for its negative effects, especially for the conditions arising from occupational hazards after prolonged exposure [2]–[4]. However, in recent years, vibration has been increasingly investigated for its positive effects. Researchers have studied the use of vibration stimulation for increasing muscle strength, muscle power, body balance and bone remodeling [5],[6],[7], [8],[9]. Consequently, given the potential benefits of vibration stimulation, it has been suggested for specific applications ranging from sports to therapeutics to rehabilitation [10], [11].

Two main types of vibration exercise modalities have been identified, whole body vibration (WBV) and upper limb vibration (ULV). WBV is delivered generally through the lower limbs with the user typically standing in a half squat position on a vibrating platform. ULV vibration devices deliver vibration stimulation to the hand and arm and can consist of vibrating dumbbells which the user grasps tightly to receive the stimulations. Both of these vibration modalities deliver stimulation indirectly, through the limbs, whereas most of our understanding about the body’s neurophysiological responses to vibration stimulation is based on earlier diagnostic studies which delivered the vibration directly to specific muscles and tendons [12], [13].

While the potential beneficial effects on muscle and bone form and functions are recognized now in various populations [14]–[16], a lack of consensus exists on the biological mechanisms responsible for such adaptations. One suggestion is that the enhanced neuromuscular activation found during WBV [16], [17] can be one of the main mechanisms inducing improvements in skeletal muscle function. For this reason, it has been suggested that WBV can induce adaptations similar to resistance training [16]–[19].

Direct vibration stimulation has been shown to enhance muscle spindle activity resulting in excitatory response of the primary and secondary endings [1], [12], the excitatory response being known as tonic vibration reflex (TVR) [13], [20]. It has also been observed that the TVR response is influenced by the vibration location, the initial length of muscle i.e. pre-stretch and the vibration frequency and amplitude [21], [22].

Another theory, proposed to explain the increased neuromuscular response under indirect vibration is muscle tuning [23]. Muscle tuning theory suggests that increased muscle contraction during vibration could lead to higher neuromuscular response [24]. Recent work has suggested some degree of a temporary sustained enhancement of corticospinal excitability concomitant with spinal inhibition acutely after WBV [25], suggesting that central aspects should not be discounted.

Considering the effect of muscle length, contraction and vibration frequency as well as amplitude in grading the neuromuscular responses to vibration stimulation, recently authors have investigated neuromuscular activation in upper limbs under vibration stimulation with isometric contractions superimposed [26], [27]. In the lower limbs, limited studies exist, but it has been shown that additional load determines an increase in EMG activity in the target muscles [28].

To the best of the authors’ knowledge, this is the first study to investigate the effects of graded isometric contractions superimposed on WBV exercise. This is also the first attempt to understand collectively, the effects of different frequencies and amplitudes of vibration stimulation when graded levels of contraction are superimposed on WBV. Another unique aspect of this study is the investigation of agonist/antagonist co-activation in the lower limbs under graded isometric exercise superimposed on WBV.

The purpose of this study was to quantify and analyze the effects of variations in the vibration parameters and contraction levels on the neuromuscular responses to isometric exercise superimposed on WBV stimulation.

To investigate the above mentioned novel aspects, we hypothesized that compared to the Control (C) condition:

1. The Vibration intervention i.e. vibration (V) condition would significantly enhance neuromuscular activation (EMGrms) in the Vastus Lateralis (VL), Vastus Medialis (VM) and Biceps Femoris (BF) muscles
2. This neuromuscular activation (EMGrms) would vary significantly between and would depend upon the vibration frequency, amplitude and isometric contraction level
3. The V condition would also significantly increase the agonist-antagonist co-activation
4. The V condition would significantly increase the peripheral fatigue indices (EMG-Mean Frequency (MEF) and Median Frequency (MDF)) in the VL, VM and BF muscles
5. These peripheral fatigue indices (EMG-MEF and MDF) would vary significantly and depend upon the vibration frequency, amplitude and isometric contraction level

## Methods

### Participants

6 female and 6 male (Age 28 years ± 7.24, Height 173 cm± 13.04, Weight 73.16 Kg± 11.19) healthy volunteer participants were recruited through the University of Aberdeen, Biomedical Engineering laboratory. The level of physical training of the participants varied from sedentary to amateur athlete. Written and informed consent was signed by each volunteer. Exclusion criteria included a history of back pain, acute inflammations in the pelvis and/or lower extremity, acute thrombosis, bone tumors, fresh fracture, fresh implants, gallstones, kidney or bladder stones, any disease of the spine, peripheral vascular disease, or pregnancy.

### Experimental setup

Trials were performed with the set-up previously reported in [29], [30]. A leg press machine was converted into a WBV device which allowed the user to apply varying levels of isometric contractions while receiving vibration stimulation in a seated position (Refer to Figures 1 and 2 for a photograph of the device and a schematic of the experimental set-up and instrumentation [31]). The leg press machine was fitted with two contra-rotating motors (Vibratechniques Ltd., UK; model: MVSI-S90) attached to a spring mounted vibration plate. The vibration plate was attached directly to the foot plate, against which user pushed to receive the vibrations. This led to the sinusoidal motion of the foot plate in the sagittal direction, towards and away from the user.

**Figure 1:**
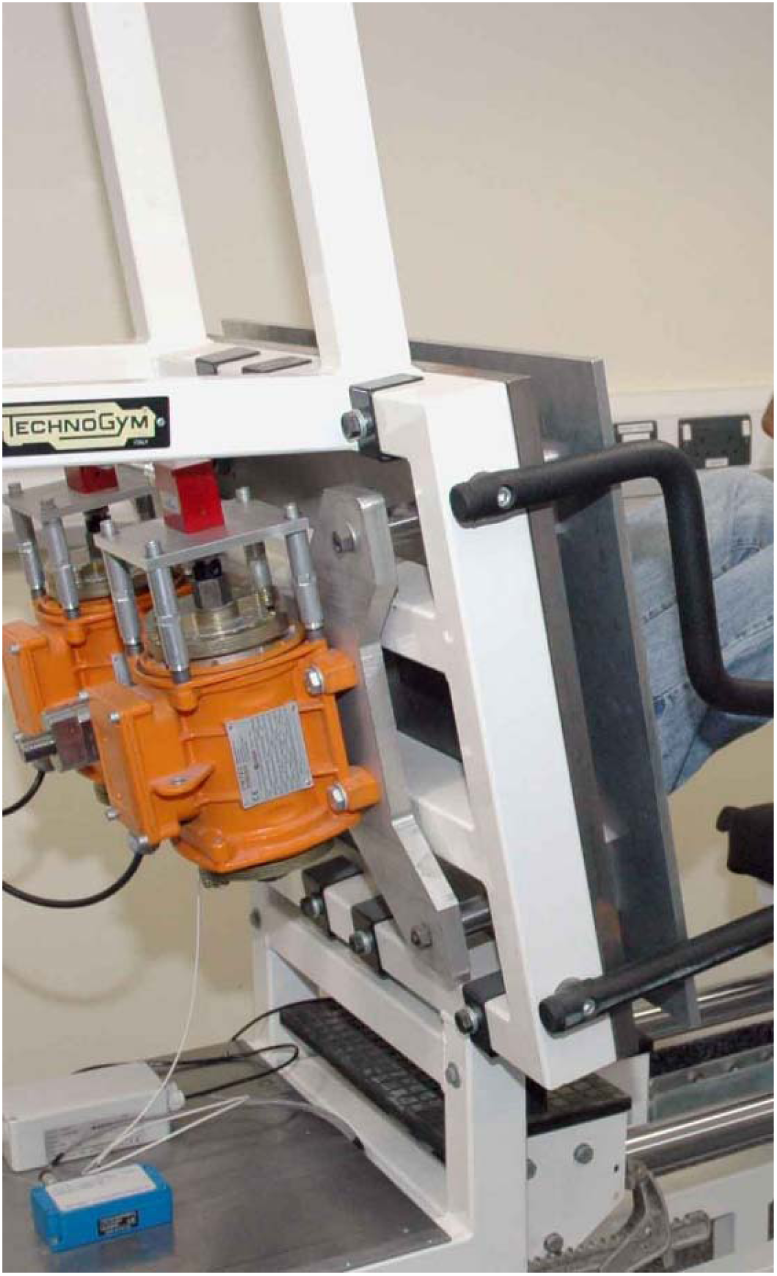
Photograph showing the arrangement of various plates delivering vibration of the motors to the user via footplate.

**Figure 2:**
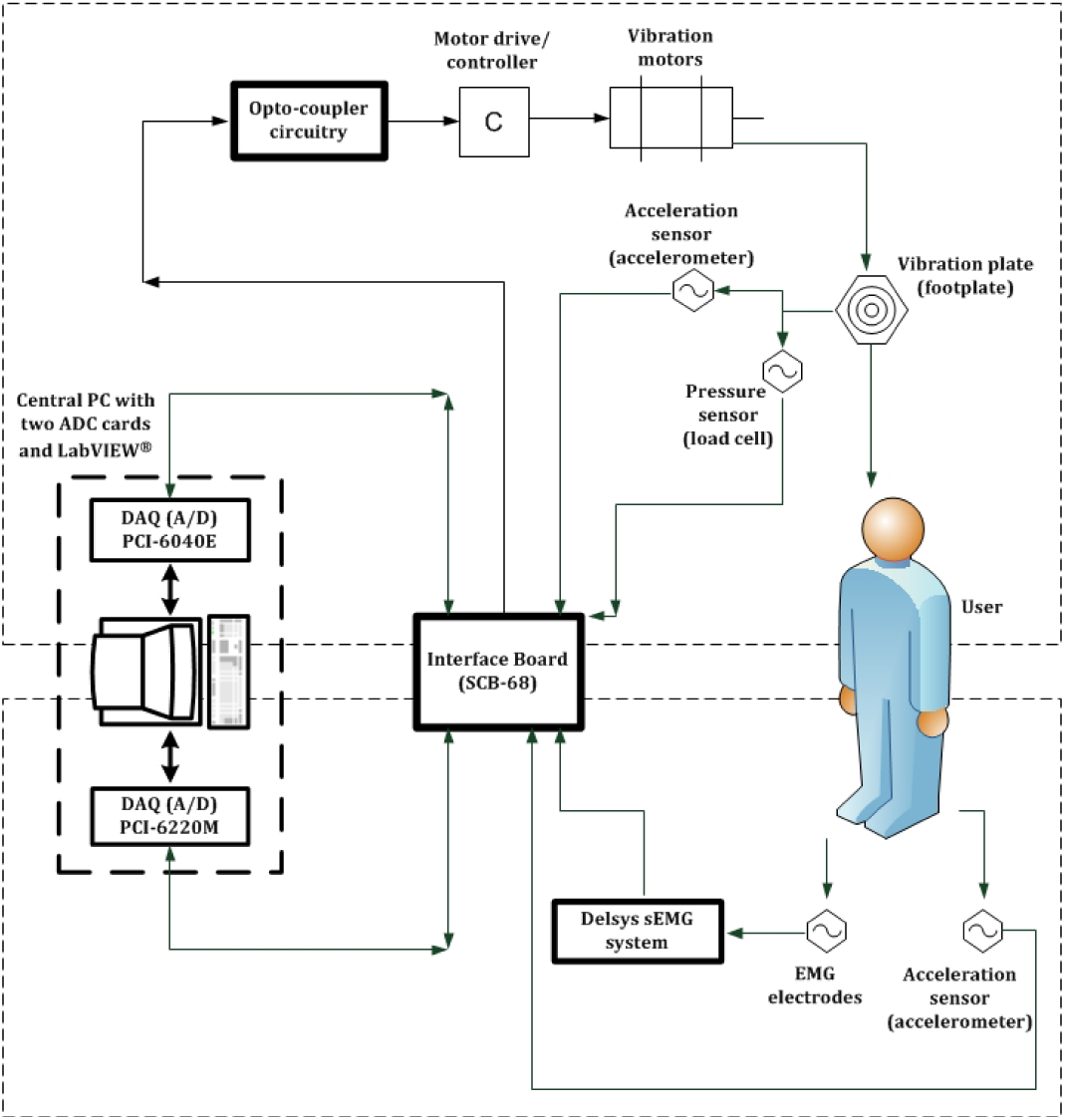
Schematic diagram showing operation of the complete WBV system, the direction of the arrow represents the flow of the signal.

An accelerometer (Kistler Instrument Corp.; model: KShear-8704B25) attached to the vibration plate sensed the real time acceleration of the vibrating plate. A pancake type load cell (Procter & Chester Measurements Ltd., UK; model: BD-PLC-C) sandwiched between vibration plate and the foot plate measured the real-time force applied by the user, i.e. MVC.

The user sat on the device seat with a backrest, with his/her legs half flexed (90⍰ knee angles) and pushed against the vibrating foot plate. This posture arrangement differs from current WBV devices where the user stands on a vibrating platform. While exercising, the knee angle was kept at 90° and was continuously monitored with a goniometer. The position of the seat and backrest could be adjusted manually with respect to the foot plate, to accommodate users of different height. This also helped to keep the knee angle of 90°. Appropriate toe and heel positions were marked on the foot plate to ensure consistency inter and intra participants. To avoid vibration damping and any variability among participants, exercises were performed bare-foot. The foot plate had a rubber platform to provide traction while exercising.

The WBV device was set up to generate sinusoidal vibrations of 30Hz or 50Hz frequencies with peak-to-peak (p-p) amplitudes of 0.5mm or 1.5mm.

### Vibration device-performance evaluation

To verify the WBV device generates specific and repeatable vibrations (i.e. 30Hz and 50Hz frequencies with peak-to-peak (p-p) amplitudes of 0.5mm and 1.5mm), a simple procedure was followed. An input voltage was delivered to the vibration device from the central PC and the corresponding vibration characteristics generated were recorded. This procedure was repeated multiple times to make sure those vibration characteristics values were consistent and hence reliably repeatable. The experimental set-up to generate the vibrations is described in schematic in Figure 3. The vibration characteristics were recorded by the accelerometer situated on the vibration plate. These signals were recorded, observed and analysed on an oscilloscope. The signal to the oscilloscope from the accelerometer was low pass filtered with a cut-off frequency 80Hz. This low-pass filtering made sure that the frequency range of interest (30Hz and 50Hz) was recorded and analysed.

**Figure 3:**
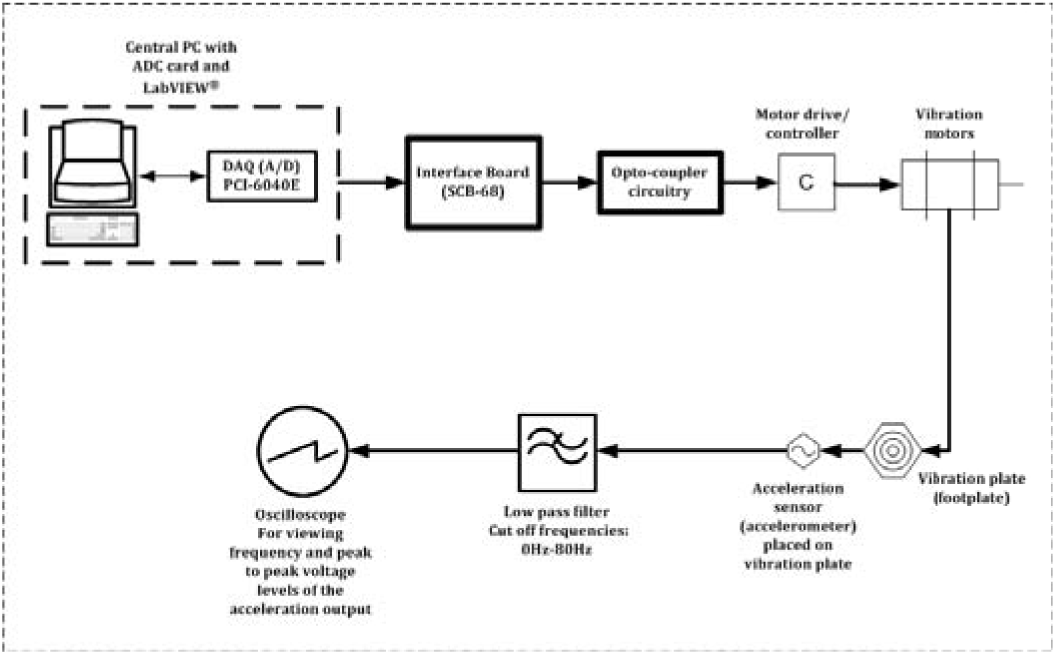
Schematic showing experimental set-up used for recording vibration characteristics to verify and evaluate the performance of the WBV system.

One set of representative data recording values are presented below in the Figure 4.

**Figure 4:**
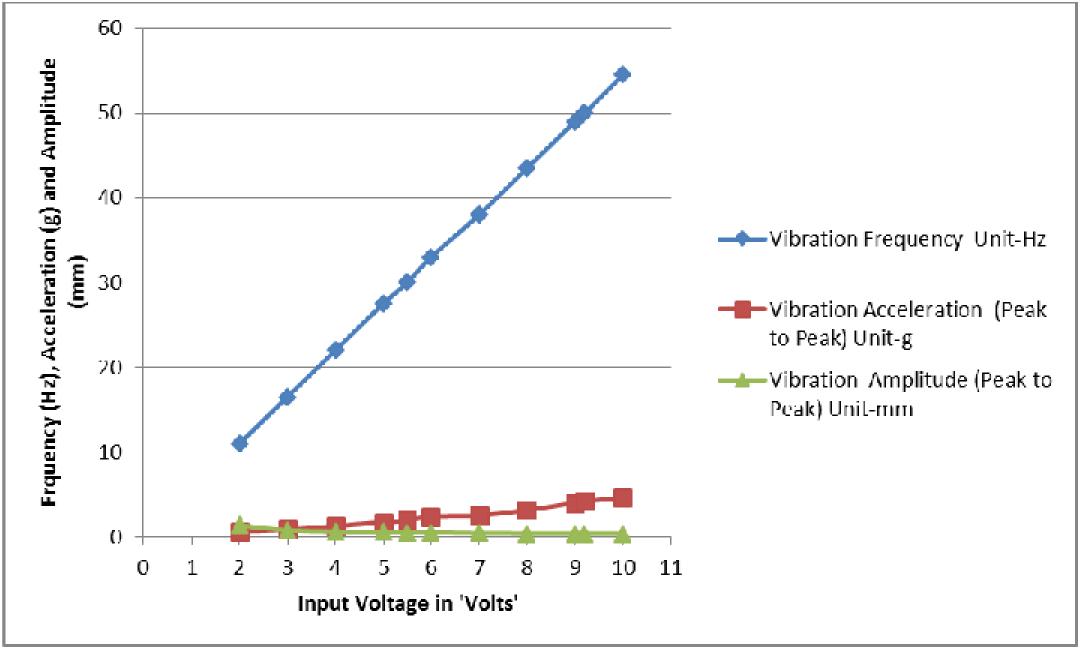
Representative data showing one et of frequency, acceleration and amplitude levels obtained from the WBV (plate) actuator against corresponding input voltages.

### Study design

A randomized cross over design was used to carry out the exercise interventions. As a cross over experiment, each participant was randomly allocated one of the several possible sequences of interventions. 25 integers were assigned to the 25 different interventions to be performed. A ‘random sequence’ i.e. a sequence consisting of 25 integers ≅ 25 interventions was assigned to each participant to provide the interventions in a random order. The MATLAB function ‘randperm’ was used to generate these random sequences.

The 25 interventions consisted of the exercise conditions shown in Table 1.

**Table 1:** Characteristics of the 25 interventions and the integers assigned to them. Control refers to ‘no vibration’ and MVC refers to Maximum Voluntary Contraction.

| Integer assigned | Characteristics of the corresponding intervention/treatment performed |
| --- | --- |
| 1 | Control- 20% of MVC |
| 2 | Control- 40% of MVC |
| 3 | Control- 60% of MVC |
| 4 | Control- 80% of MVC |
| 5 | Control- 100% of MVC |
| 6 | 30Hz+0.5mm+20% of MVC |
| 7 | 30Hz+0.5mm+40% of MVC |
| 8 | 30Hz+0.5mm+60% of MVC |
| 9 | 30Hz+0.5mm+80% of MVC |
| 10 | 30Hz+0.5mm+100% of MVC |
| 11 | 30Hz+1.5mm+20% of MVC |
| 12 | 30Hz+1.5mm+40% of MVC |
| 13 | 30Hz+1.5mm+60% of MVC |
| 14 | 30Hz+1.5mm+80% of MVC |
| 15 | 30Hz+1.5mm+100% of MVC |
| 16 | 50Hz+0.5mm+20% of MVC |
| 17 | 50Hz+0.5mm+40% of MVC |
| 18 | 50Hz+0.5mm+60% of MVC |
| 19 | 50Hz+0.5mm+80% of MVC |
| 20 | 50Hz+0.5mm+100% of MVC |
| 21 | 50Hz+1.5mm+20% of MVC |
| 22 | 50Hz+1.5mm+40% of MVC |
| 23 | 50Hz+1.5mm+60% of MVC |
| 24 | 50Hz+1.5mm+80% of MVC |
| 25 | 50Hz+1.5mm+100% of MVC |

The protocol is outlined in Figure 5. In the first visit each participant was familiarized with the WBV device. Then the participant performed an isometric leg press exercise of various intensities against the WBV device foot-plate with the knees flexed at 90° as a warm up. After this initial warm-up and familiarization, the maximal voluntary contraction (MVC) was established for each participant. For this, the participant performed maximal efforts for 40 seconds. This procedure was undertaken 3 times with each effort separated by a 5 minute period of rest. The average of the three efforts was used as the baseline MVC value for that participant.

**Figure 5:**
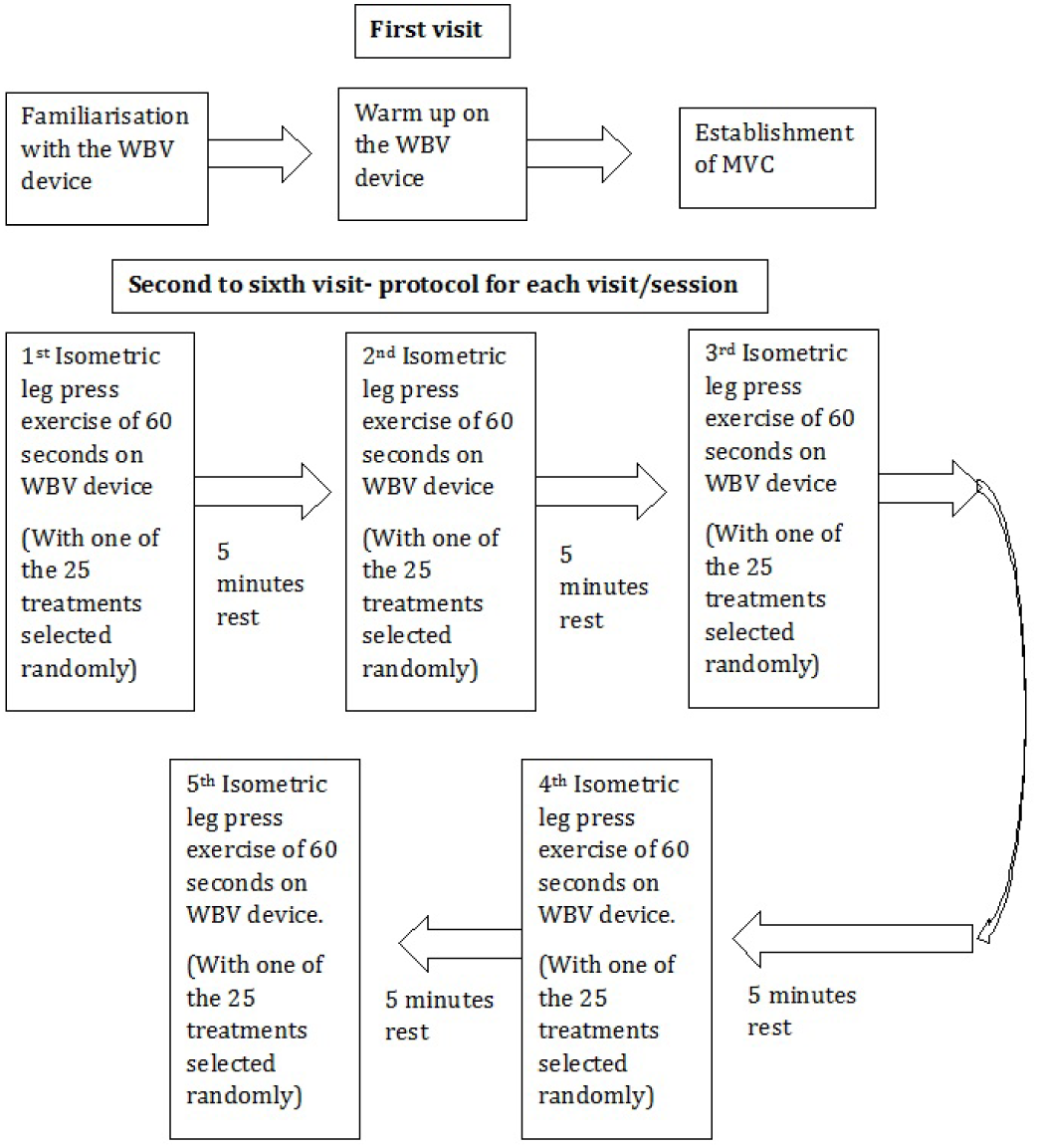
Schematic describing the arrangements/research protocol of the experiment

In the second to sixth visits the participants went through the vibration exercise intervention outlined in Table 1 with the randomized cross-over design. The vibration interventions consisted of an isometric leg press exercise pushing against the WBV device foot-plate with the knees flexed at 90° at the target force, for 60 seconds. 5 minutes rest was allowed between any two consecutive interventions/measurements. 5 interventions were carried out per session with the total of 25 measurements taken in the five sessions.

During each measurement, the neuromuscular activation of the designated muscles was recorded in the form of surface electromyographic (sEMG) response and stored for analysis along with the vibration characteristics and the force production details. The vibration being delivered was continuously monitored and recorded. Vibrations transmitted to the participant were recorded and stored by a tri-axial accelerometer (Analog Devices Inc.; model: ADXL-330) attached to the lower limb of the participant. Real time graphical and numerical representations of the vibration characteristics as well as the force levels produced by the participant were available on the main computer.

All the procedures were non-invasive. Participant wore shorts to facilitate sensor placement on the lower limbs.

### Instructions to the participants

Participants were asked to maintain consistency in their foot positions and knee angles. Both of these variables were measured continuously throughout the tests. Participants received both verbal and visual (real time graphical values on the PC) feedback to assist them in maintaining a constant force level.

Isometric contraction with the required postural conditions was practiced with and without vibrations before the actual trials until the participants became familiar with the test conditions. Trials were repeated if postural conditions changed from the required position.

### Fatigue and Safety

A minimum of 72 hours of recovery time was allowed between any two testing sessions to avoid any residue of fatigue and/or delayed onset of muscle soreness. Also, a general log of each participant’s daily physical activity excluding the trials was kept, e.g. any form of regular/irregular physical exercise like running, strength or resistance training etc. This was done to ensure that the participant did not undergo the WBV stimulation trials immediately after finishing their regular exercise. At least 72 hours of time gap was allowed between the regular physical exercise and WBV trials, to avoid any effect of muscle fatigue.

Participants were encouraged to report any pain and/or discomfort, during or after the trials, to the test operator. Apart from the feeling of exertion during the exercises performed at an individual’s peak capacity, no adverse effects were reported by the participants during or after the trials.

An emergency stop button to halt the vibration delivery was located near the WBV device seat. Participants were advised to make use of the button in case they felt unsafe or were in pain. None of the participants used the emergency stop button during the trials.

### EMG measurements and processing

sEMG was recorded from the VL, VM, and BF during all exercise conditions according to recommendations reported in the literature [32]. Active bipolar electrodes (DelSys, Inc.; model: DE 2.1) were aligned with the muscle fiber direction and placed between the tendon and the muscle belly. To minimize the impedance and to ensure a proper contact, the skin was shaved as necessary, lightly abraded and cleaned with 70% isopropyl alcohol. The reference electrode was placed on an electrically inactive area of the lumbar spine (the anterior superior iliac spine). To ensure consistency in the placement of the sEMG electrodes between the sessions, electrode locations were marked with a skin marker and kept throughout the entire duration of tests, i.e. from the first MVC measurement visit to the last visit. The sEMG electrodes and cables were secured to subject’s skin with medical tape. Active grounding and shielding of the cables was carried out to minimize electromagnetic inference [33]. The sEMG signals were sampled at 1000Hz, amplified with a gain of 1000 and analogue filtered for a 20-450Hz band pass with DelSys hardware (DelSys, Inc.; model: Bagnoli-4). Data acquisition was performed through a 16 bit data acquisition card (National Instruments Corp.; model: PCI-6220M) and EMGWorks (DelSys, Inc.) software.

Subsequent data processing and analysis was performed with custom written MATLAB code (The Mathworks, Inc.; version 8) routine. Any baseline offset of the sEMG data was removed by subtracting the mean.

The root mean square (RMS i.e. EMGrms) was used to estimate the neuromuscular activation. The RMS was calculated using the moving window technique. Initially the RMS was calculated for each window, and then the RMS for the entire data length was obtained by averaging the individual RMS values of each window. Exactly same RMS windowing characteristics were employed to obtain the MVC EMGrms values as well as EMGrms for the C and V conditions.

The Mean Frequency (MEF) and Power Spectral Density (PSD) of the sEMG data were also obtained. The MEF was used as an indicator of muscle fatigue. These spectral estimators were also derived by moving window technique.

For both amplitude and spectral estimation, the hamming window with a length of 1 second and no overlap was used. It has been shown that the choice of the window does not have a critical bearing on the spectral estimators like MEF and PSD [34]. Further, for isometric, constant force and fatiguing contractions, the signal is regarded as stationary for epoch/window duration of 1 to 2 s. Previous studies suggest that epoch durations between 500 ms to 1 s provide better spectral estimation [34]–[36]. Also, it has been shown that window overlapping does not provide any significant benefits [34]. Based on these recommendations, the window length was kept to 1 s without any overlap.

### Line Artifact Removal

The power spectral analysis of the sEMG revealed peaks coinciding with 50Hz and to a lesser degree with 30Hz.

Some authors have filtered the peaks in sEMG spectra coinciding with the vibration stimulation frequencies assuming them to be motion artifacts [37]. However it is still unclear whether the spectral peaks correlating with the stimulation frequencies are in fact motion artifacts [36] or stretch reflexes [18]. Recent evidence suggests that these peaks can indeed be stretch reflexes [38]. Considering the present ambiguity about the existence of motion artifacts and increasing evidence towards the presence of stretch reflex [18], [38], only the spectra exhibiting the largest power and hence potential to skew the results were removed. The largest spectra were found to be at 50Hz irrespective of the stimulation frequency of 30Hz or 50Hz. Hence, a Butterworth notch filter (10 ^th^order, cut-off frequencies 49.5-50.5Hz) was employed to remove the components at this frequency.

### Statistical analysis

Normalization was performed by dividing the EMGrms of the entire section of the data value to be normalized by the maximum value obtained from the MVC effort of each participant. To identify whether the EMGrms values differ significantly between the effort levels and between the control and vibration conditions, a 2 way ANOVA test was employed. The 2 interventions (control and vibration) and 5 intensities (effort levels) were used to compare between the EMGrms of different effort levels as well as between the control and each vibration condition one at a time. Alpha was set at 0.05. In each case, a significant difference was defined for a computed p-value ≤ 0.05. Paired student t-tests (one tail, different variance) were employed at each effort level to compare the sEMG responses between the C and V conditions and to establish the significance level (P value) of the deviations from the means. The distribution of grouped data was assessed for normality using the Lilliefors test with a significance detection level of ≤ 0.05. Statistical analysis was carried out using the SigmaPlot statistical software package (Systat Software Inc.; Version SigmaPlot 12).

## Results

### Overall Effects of Vibration on EMG Amplitude

For the VL and BF muscles, at all contraction levels, isometric contraction superimposed on vibration stimulation produced higher mean EMGrms activity than isometric contraction (control) alone. However the VM did not show any increases in neuromuscular activity under vibration conditions, instead its mean EMGrms values were similar to the control condition and in some cases lower.

As a prime mover/agonist in the leg press exercise, the VL displayed higher EMG activity than the control condition. The percentage increase in mean EMGrms values with vibration was highly variable depending on the frequency, amplitude and contraction level and ranged from 5% to 165%. The EMGrms data for the various cases are shown in Figures 8-11.

**Figure 8:**
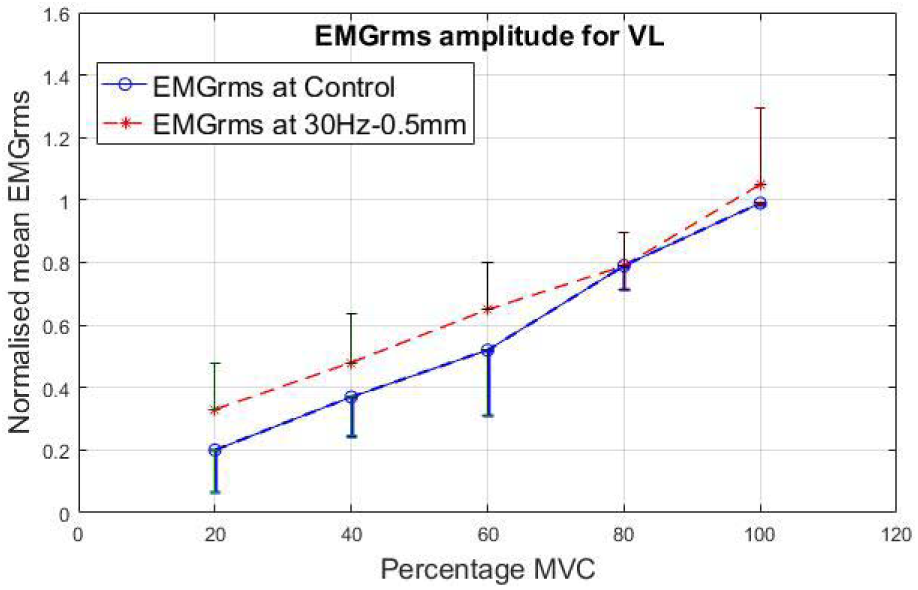
Normalised mean EMGrms values for VL at 20, 40, 60, 80 and 100% MVC under 30Hz-0.5mm V against C (no vibration) condition.

**Figure 9:**
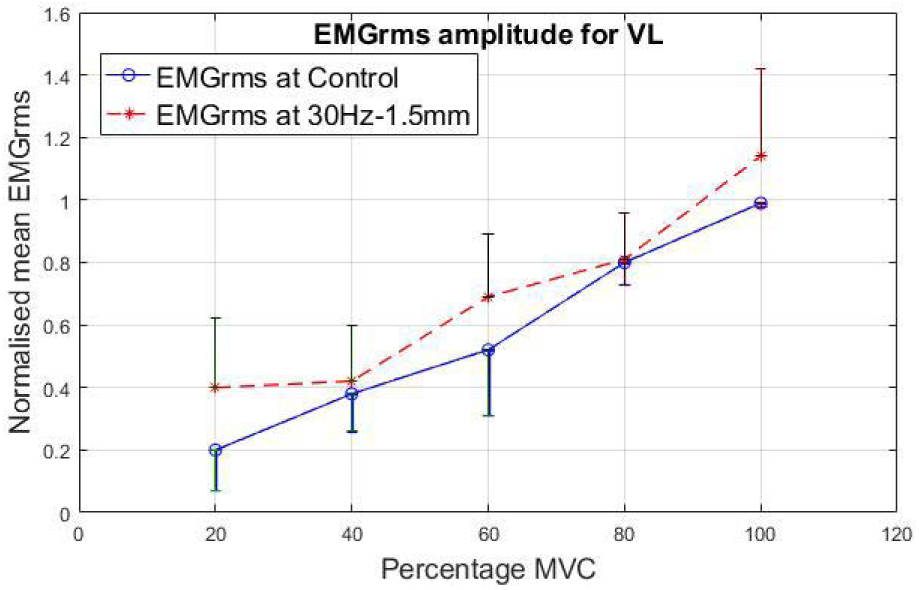
Normalised mean EMGrms values for VL at 20, 40, 60, 80 and 100% MVC under 30Hz-1.5mm V against C.

**Figure 10:**
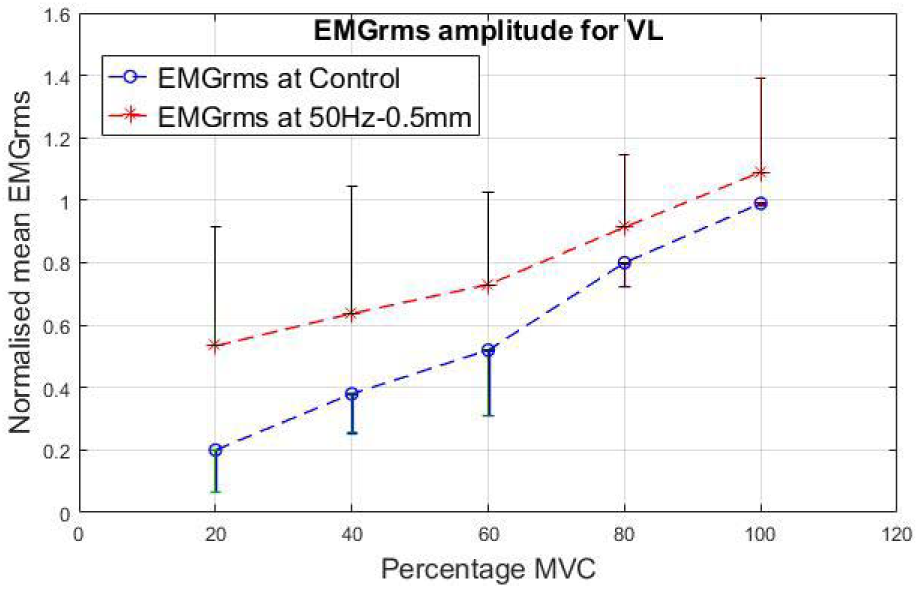
Normalised mean EMGrms values for VL at 20, 40, 60, 80 and 100% MVC under 50Hz-0.5mm V against C.

**Figure 11:**
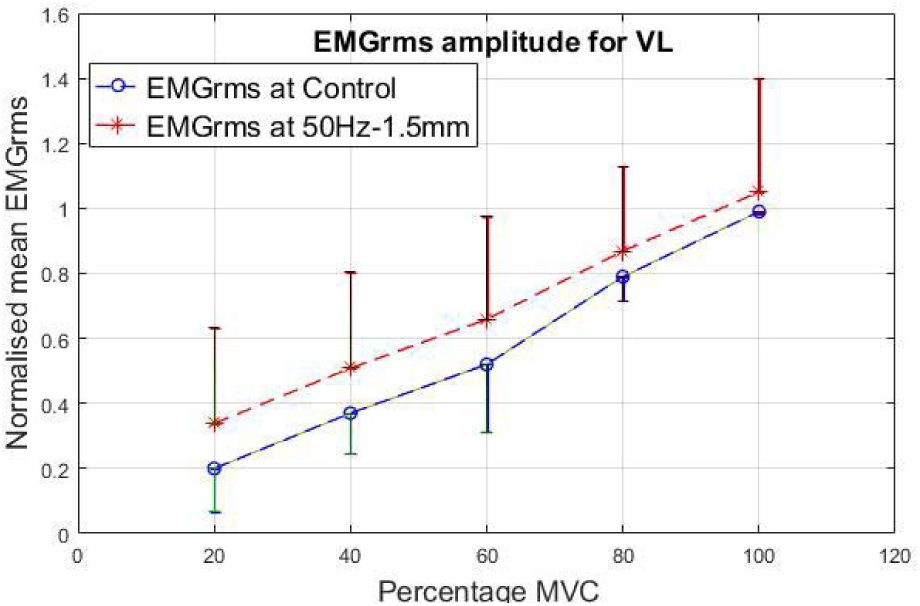
Normalised mean EMGrms values for VL at 20, 40, 60, 80 and 100% MVC under 50Hz-1.5mm V against C.

As an antagonist, the BF seemed highly active and showed higher levels of EMG activity under vibration compared to the control condition. Similar to the VL, the percentage increase in mean EMGrms values of BF was highly variable and depended on the frequency, amplitude and muscle contraction level. Compared to the control, the BF’s mean EMGrms increase ranged from 28% to 206%. The EMGrms data for the various cases are shown in Figures 12-15.

**Figure 12:**
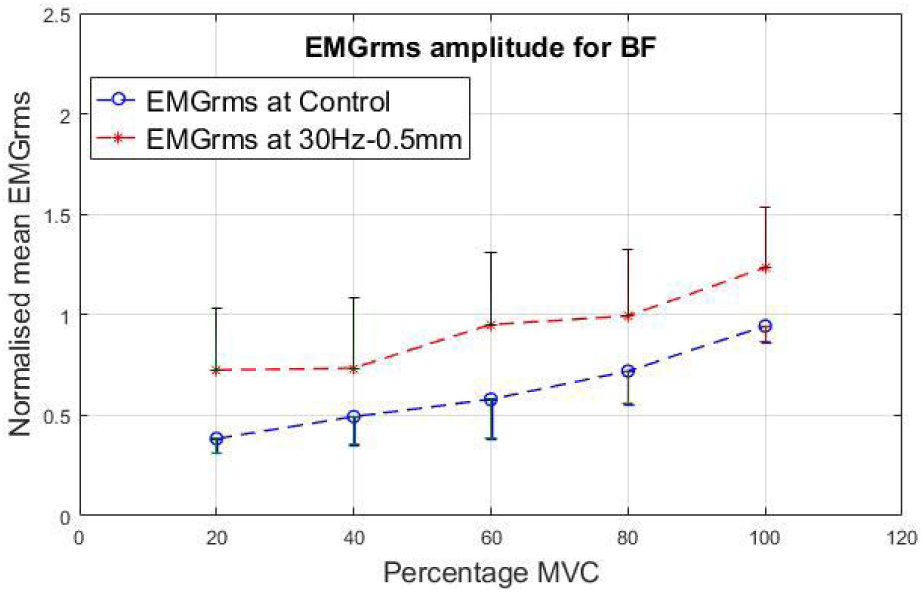
Normalised mean EMGrms values for BF at 20, 40, 60, 80 and 100% MVC under 30Hz-0.5mm V against C.

**Figure 13:**
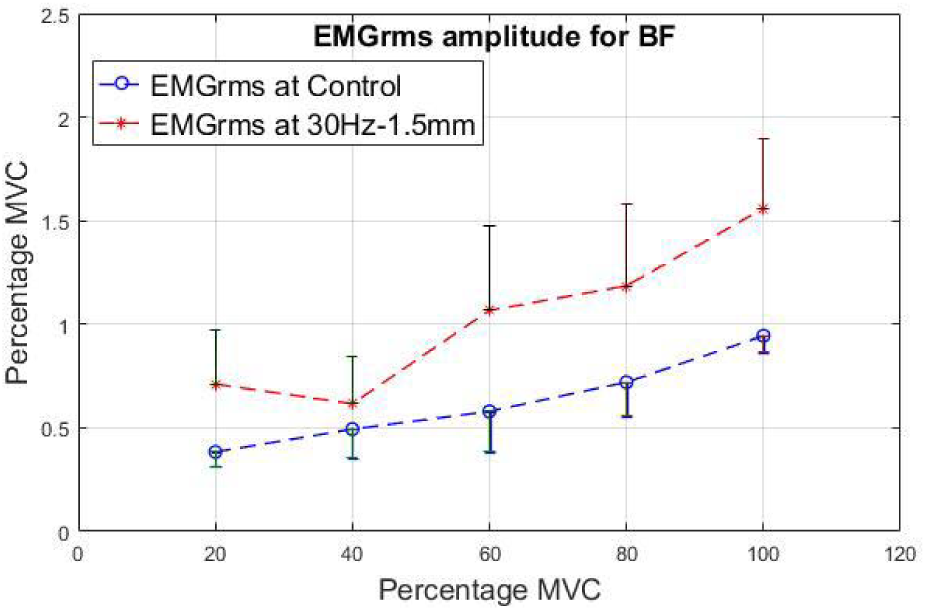
Normalised mean EMGrms values for BF at 20, 40, 60, 80 and 100% MVC under 30Hz-1.5mm V against C.

**Figure 14:**
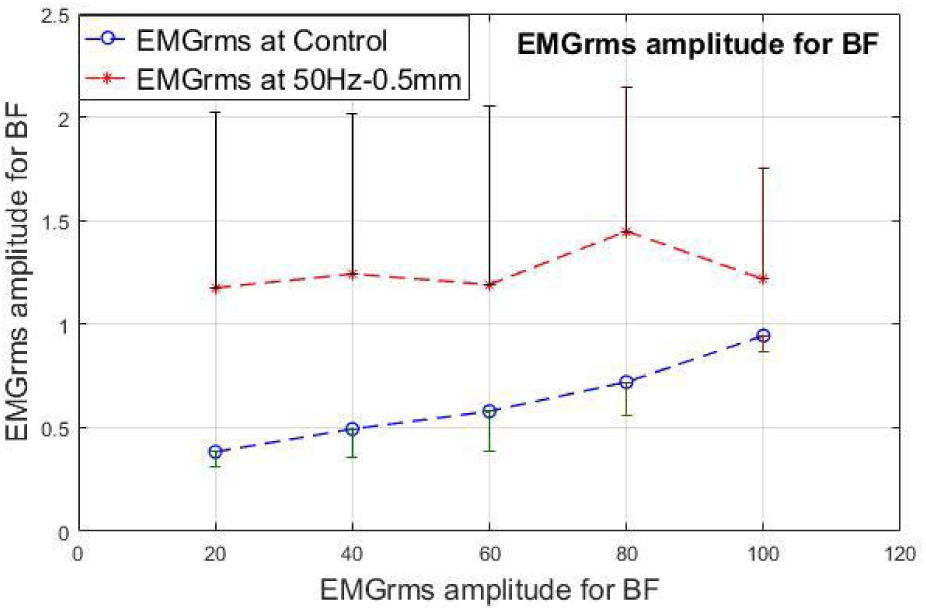
Normalised mean EMGrms values for BF at 20, 40, 60, 80 and 100% MVC under 50Hz-0.5mm V against C.

**Figure 15:**
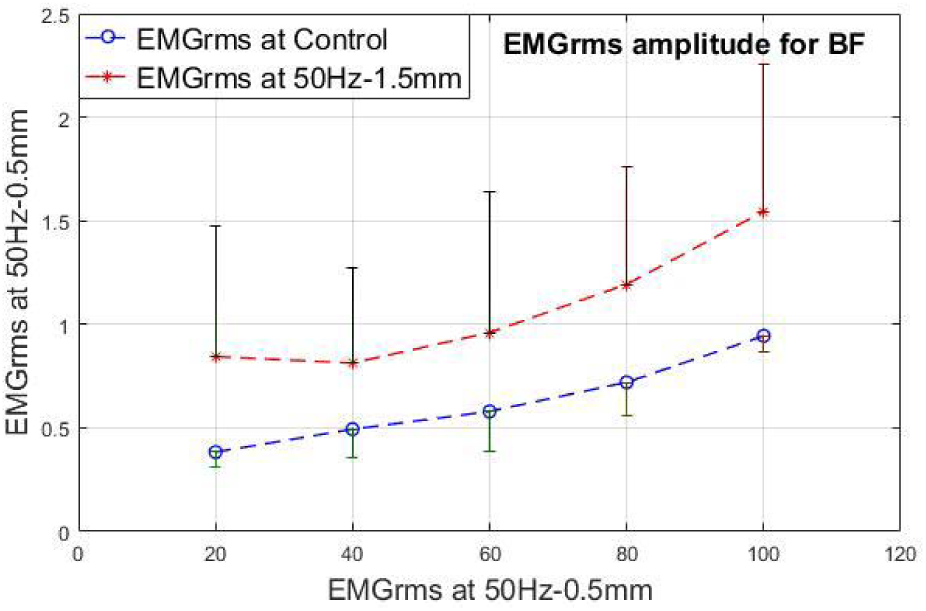
Normalised mean EMGrms values for BF at 20, 40, 60, 80 and 100% MVC under 50Hz-1.5mm V against C.

### The Effects of Frequencies 30Hz and 50Hz and Amplitudes 0.5mm and 1.5mm

Among the four combinations of the vibration variables investigated, 50Hz-0.5mm stimulation induced the largest neuromuscular activity in the VL and BF muscles with the highest increases of 165% and 206% in mean EMGrms values respectively (Table 2 and Figures 8-15).

**Table 2:** Percentage increase or variation in mean EMGrms values in comparison with respective controls. Values in bold represent statistically significant increase compared to C with P value ≤ 0.05.

| Vibration treatment condition (Frequency= 30/ 50Hz, amplitude= 0.5/1.5mm, Force level = 20/40/60/80/100% of MVC) | Vibration Treatment effect in a Muscle Group (% increase over respective control condition) |  |  |
| --- | --- | --- | --- |
|  | VL | VM | BF |
| 30Hz_0.5mm_20 | 66.85* | 25.72 | 89.22* |
| 30Hz_0.5mm_40 | 28.81 | -5.17 | 48.98* |
| 30Hz_0.5mm_60 | 26.43* | 10.67 | 64.33* |
| 30Hz_0.5mm_80 | -0.14 | -10.69 | 37.99* |
| 30Hz_0.5mm_100 | 5.96 | 0.50 | 30.81* |
| 30Hz_1.5mm_20 | 98.43* | 23.56 | 85.45* |
| 30Hz_1.5mm_40 | 10.11 | -13.70 | 24.87* |
| 30Hz_1.5mm_60 | 31.92* | 23.09 | 84.15* |
| 30Hz_1.5mm_80 | 2.01 | -4.377 | 64.55* |
| 30Hz_1.5mm_100 | 14.75 | 11.010 | 65.12* |
| 50Hz_0.5mm_20 | 165.44* | 15.24 | 206.50* |
| 50Hz_0.5mm_40 | 68.97* | -9.44 | 151.68* |
| 50Hz_0.5mm_60 | 40.07* | 6.62 | 105.56* |
| 50Hz_0.5mm_80 | 14.82 | -8.65 | 101.05* |
| 50Hz_0.5mm_100 | 9.86 | -1.56 | 28.86 |
| 50Hz_1.5mm_20 | 68.51 | 34.35 | 120.04* |
| 50Hz_1.5mm_40 | 34.83 | -9.02 | 64.74* |
| 50Hz_1.5mm_60 | 26.36 | 9.68 | 65.70* |
| 50Hz_1.5mm_80 | 9.18 | -2.73 | 65.64* |
| 50Hz_1.5mm_100 | 5.83 | -1.13 | 63.54* |

Interestingly, the VL did not display significantly higher EMG amplitude values under the higher level stimulation of 50Hz-1.5mm (Figure 11), whereas it recorded significantly higher (P < 0.05) EMG activity under 30Hz-1.5mm at both 20% and 60% of MVC effort (Figure 9). However, for the same vibration input i.e. 30Hz-1.5mm, the BF did not respond well compared to 50Hz-1.5mm, for which the BF generated significantly higher (P < 0.05) EMG activity at all effort levels with 63 to 120% increases in mean EMGrms values (Figures 13 and 15). Thus, apart from the 50Hz-0.5mm stimulation there was no clear combination of vibration variables which was able to generate consistently significant levels of neuromuscular response in both agonist and antagonist muscles simultaneously.

Broadly speaking, based on the percentage increases in mean EMGrms activities of the VL and BF muscles, the 50Hz-0.5mm stimulation induced the largest neuromuscular response followed by 50Hz-1.5mm and 30Hz-1.5mm (Refer Table 2). However, the EMG amplitudes under 50Hz-1.5mm and 30Hz-1.5mm stimulations were not significantly different to each other.

EMGrms Amplitude Differences between the Effort Levels and between the Control

Table 3 shows the comparison between the EMGrms of different effort levels (i.e. between 20, 40, 60, 80 and 100% of MVC) and between vibration and control condition. The values are presented for all the three muscle groups studied i.e. VL, VM and BF.

**Table 3:** 2 way ANOVA results comparing EMGrms means between effort levels and between control and vibration

| Intervention and Muscle group | Effect of treatment-effort levels (Significant difference between effort levels, P value) | Effect of treatment- V or C condition (Significant difference between V and C condition, P value) |
| --- | --- | --- |
| 30Hz-0.5mm- VL | Yes, $P < 0.001$ | Yes, $P = 0.029$ |
| 30Hz-1.5mm- VL | Yes, $P < 0.001$ | Yes, $P = 0.035$ |
| 50Hz-0.5mm- VL | Yes, $P = 0.003$ | Yes, $P = 0.010$ |
| 50Hz-1.5mm- VL | Yes, $P < 0.001$ | Yes, $P = 0.003$ |
| 30Hz-0.5mm- VM | Yes, $P < 0.001$ | No, $P = 0.966$ |
| 30Hz-1.5mm- VM | Yes, $P < 0.001$ | No, $P = 0.376$ |
| 50Hz-0.5mm- VM | Yes, $P < 0.001$ | No, $P = 0.542$ |
| 50Hz-1.5mm- VM | Yes, $P < 0.001$ | No, $P = 0.712$ |
| 30Hz-0.5mm- BF | Yes, $P < 0.001$ | Yes, $P < 0.001$ |
| 30Hz-1.5mm- BF | Yes, $P = 0.024$ | Yes, $P = 0.009$ |
| 50Hz-0.5mm- BF | Yes, $P = 0.314$ | Yes, $P = 0.003$ |
| 50Hz-1.5mm- BF | Yes, $P = 0.005$ | Yes, $P < 0.001$ |

### The Effects of Contraction Levels 20 to 100% MVC

Overall, statistically significant (P < 0.05) differences were observed between the effort levels’ mean EMGrms values. That is, as the force level increased the EMGrms values also increased significantly (Table 3). ANOVA showed significant differences between the mean EMGrms values of the effort levels in all the muscle groups. Significant differences also existed between the mean EMGrms values of all the control and vibration conditions of VL and BF muscles (Table 3).

For the VL, based on the percentage increases in mean EMGrms values, force levels of 20% to 60% of MVC seemed to induce higher neuromuscular responses than 80 and 100% of MVC efforts (Table 2). At 80% to 100% of MVC, the mean EMGrms values were similar to the control condition.

For increasing contraction levels, the VM did not display any significantly higher EMG activity for vibration compared to the control.

### Agonist-Antagonist Co-activation

Co-activation was calculated for the ratio of the EMG of the BF divided by the VL. The results are shown in Figures 16-19 for the different interventions. The EMGrms ratio of BF/VL, showed higher co-activation values with the vibration condition than the control condition except at 20% of the MVC.

**Figure 16:**
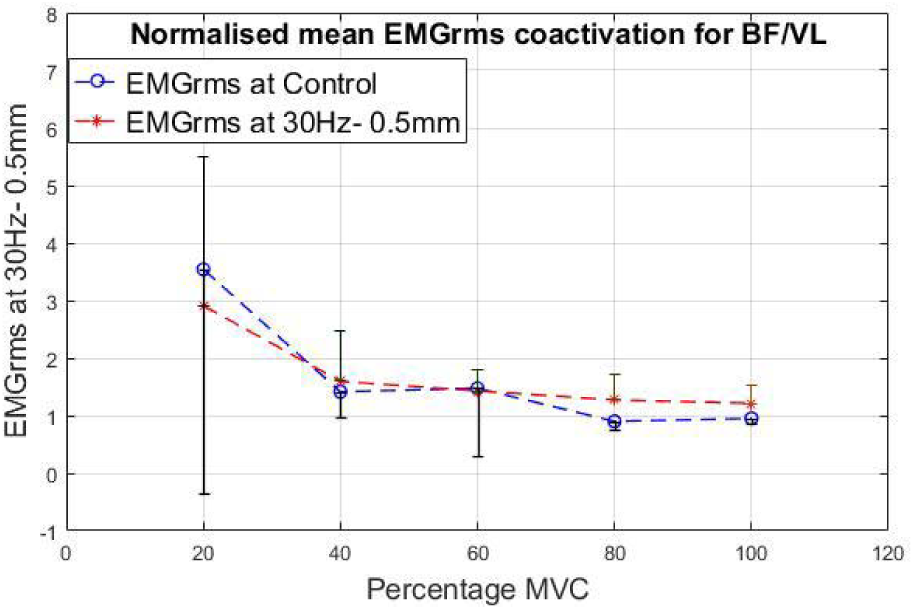
Normalised mean EMGrms co-activation values for BF over VL at 20, 40, 60, 80 and 100% MVC under 30Hz-0.5mm V against C.

**Figure 17:**
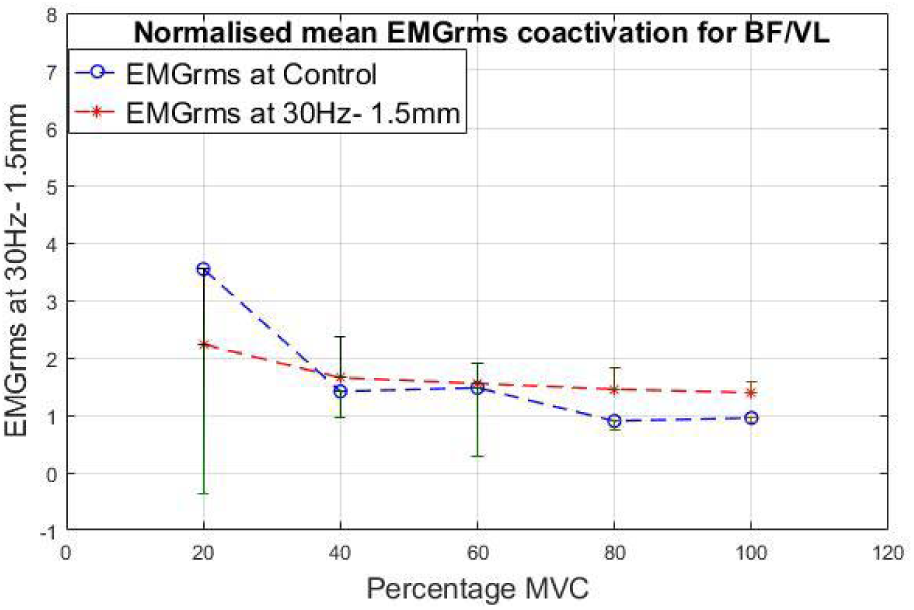
Normalised mean EMGrms co-activation values for BF over VL at 20, 40, 60, 80 and 100% MVC under 30Hz-1.5mm V against C.

**Figure 18:**
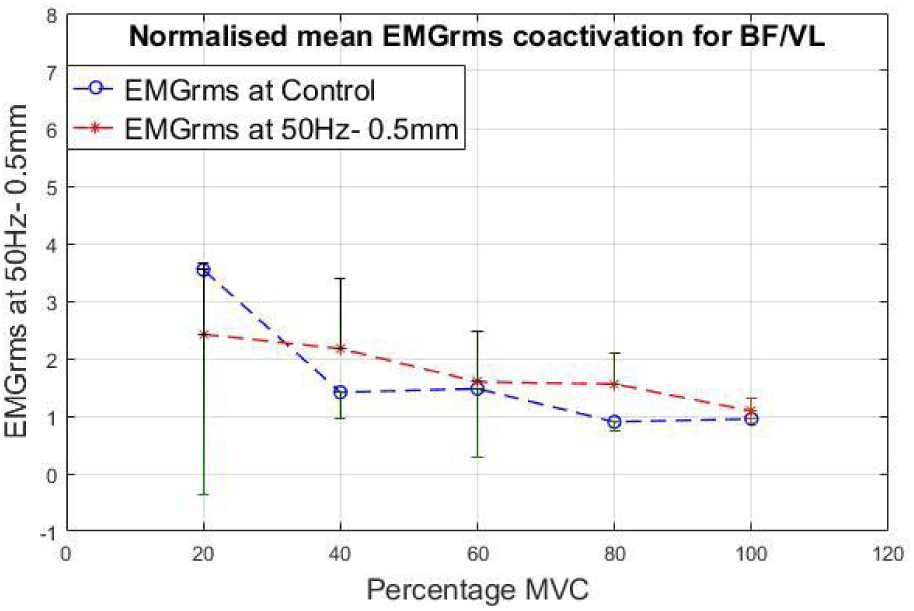
Normalised mean EMGrms co-activation values for BF over VL at 20, 40, 60, 80 and 100% MVC under 50Hz-0.5mm V against C.

**Figure 19:**
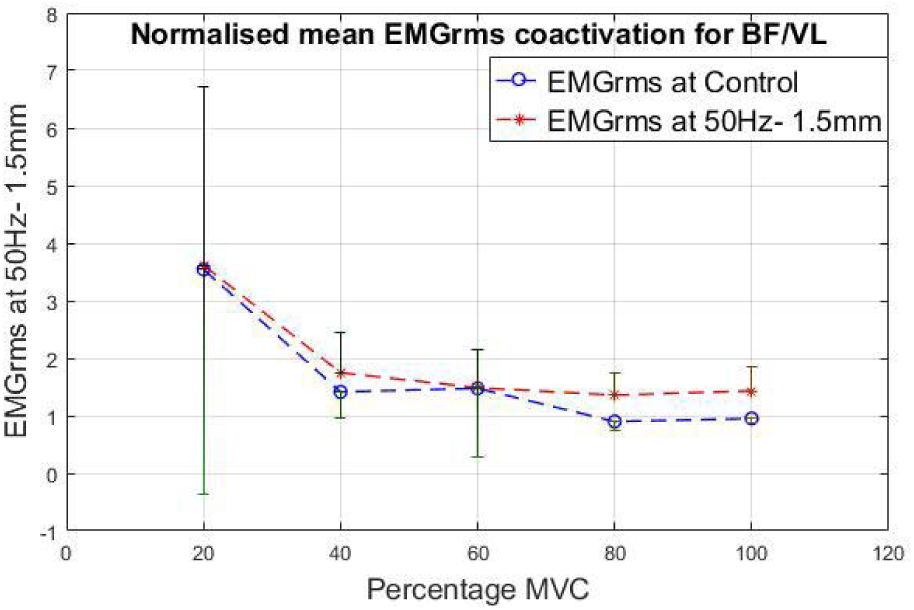
Normalised mean EMGrms co-activation values for BF over VL at 20, 40, 60, 80 and 100% MVC under 50Hz-1.5mm V against C.

As the contraction level increased, overall co-activation EMG amplitude decreased both under vibration and control conditions, with the highest co-activation amplitude being produced at 20% of MVC and the lowest at 100% of MVC. Despite an overall decrease in the co-activation amplitude with the increasing contraction, effort levels of 80% and 100% of MVC led to the most significantly (P < 0.05) higher co-activation ratios compared to the control, irrespective of the vibration condition (Figures 16-19).

50Hz-0.5mm vibration condition led to the strongest co-activation response with co-activation ratios significantly (P < 0.05) higher at 40, 80 and 100% of MVC than the control. This suggests that the higher the vibration stimulus is (i.e. 50Hz-0.5mm), the higher the co-activation required to stabilize the joint rotation during vibration. This implies 50Hz-0.5mm to be the most efficacious stimulus among the variables tested for this thesis.

For all the effort levels and vibration conditions, BF/VM co-activation was higher under the vibration condition than the control condition.

### Overall Effects of Vibration on EMG Mean Frequency Behaviour

The VL and BF show higher MEF values under all vibration conditions compared to the control. The results are shown in Figures 20-23 for the different interventions. The VM MEF values under vibration conditions did not differ much compared to the control condition’s mean frequencies.

**Figure 20:**
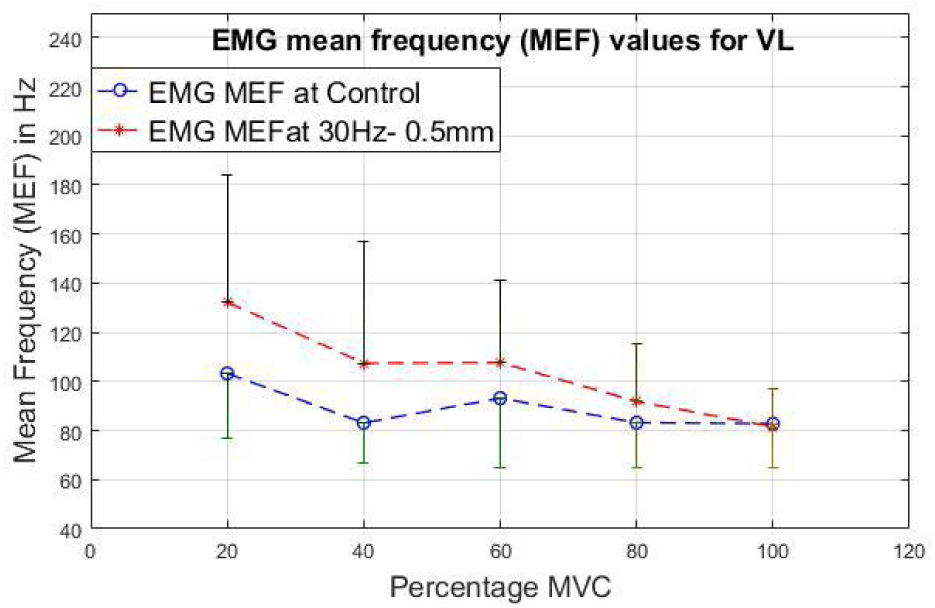
EMG mean frequency values for VL at 20, 40, 60, 80 and 100% MVC under 30Hz-0.5mm V against C.

**Figure 21:**
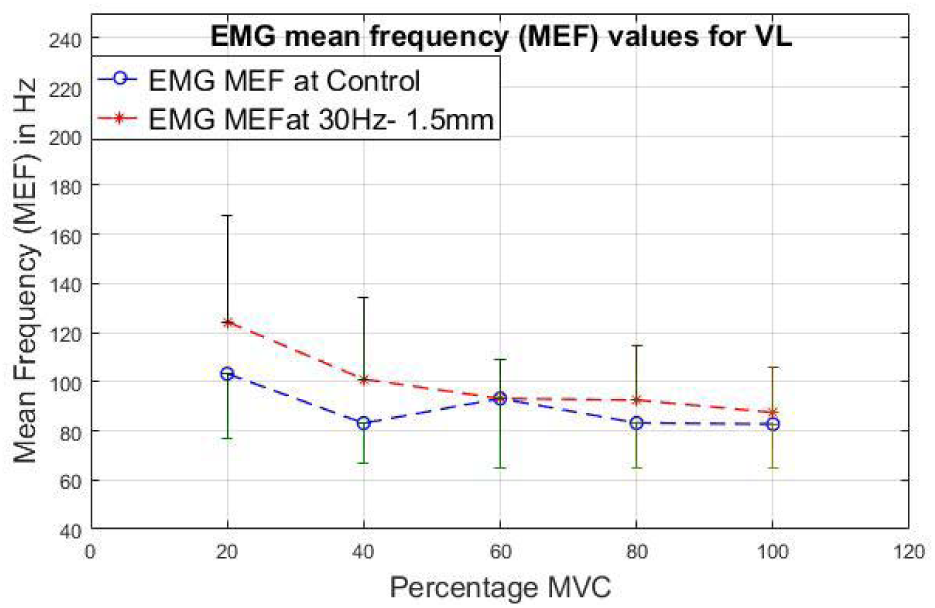
EMG mean frequency values for VL at 20, 40, 60, 80 and 100% MVC under 30Hz-1.5mm V against C.

**Figure 22:**
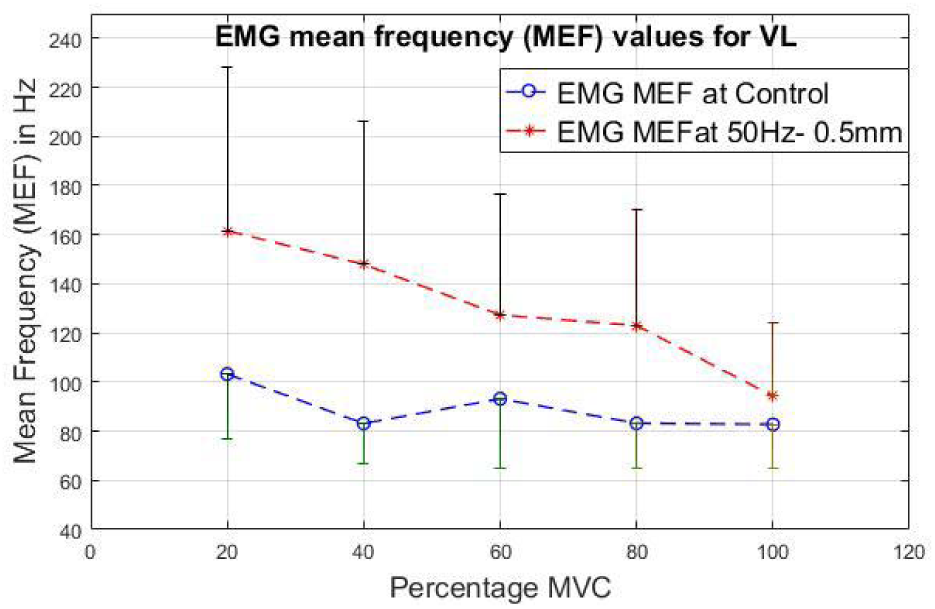
EMG mean frequency values for VL at 20, 40, 60, 80 and 100% MVC under 50Hz-0.5mm V against C.

**Figure 23:**
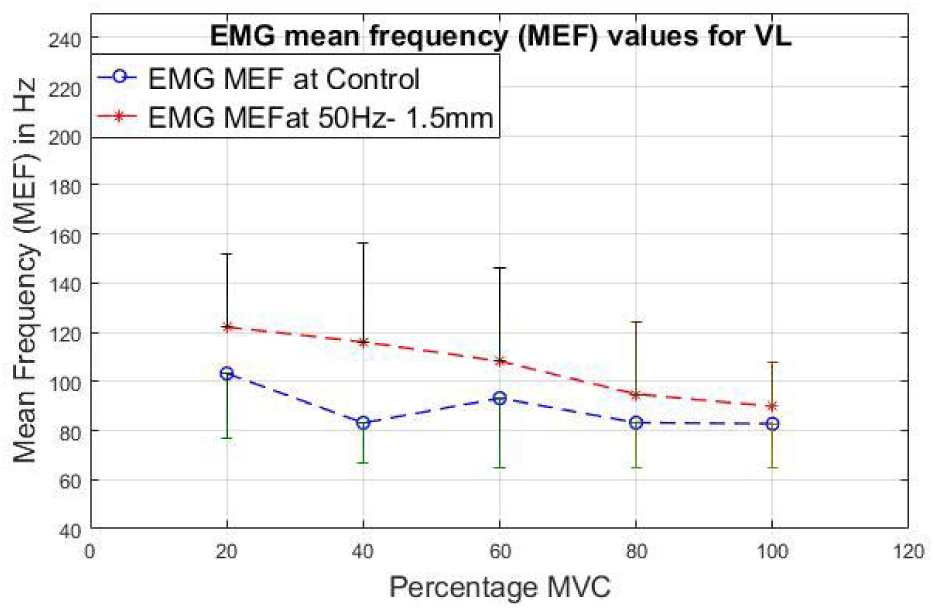
EMG mean frequency values for VL at 20, 40, 60, 80 and 100% MVC under 30Hz-1.5mm V against C.

For the VL (Figures 20-23), the lower contraction levels of 20 and 40% of MVC produce consistently the most significantly significant higher (P < 0.05) MEF values with vibration compared to control conditions.

For the BF, all contraction levels i.e. 20 to 100% of MVC produce significantly higher (P < 0.05) MEF values under specific vibration conditions compared to the control conditions. The BF display significantly higher (P < 0.05) EMG MEF values under 50Hz-0.5mm and 50Hz-1.5mm, under all contraction levels with the exception of 40% of MVC.

Although, the VM MEF values were closer to the control MEF, the VM did display higher MEF under certain vibration conditions (30Hz-1.5mm and 50Hz-0.5mm) compared to the control. However, under certain conditions its values were lower than the control’s (30Hz-0.5mm and 50Hz-1.5mm).

For both the VL and BF, the difference between the vibration and control condition MEF is larger at lower contraction levels and this difference reduces with increase in the contraction level.

## Discussion

### Effects of Vibration frequency, Vibration amplitude and Contraction levels

These results (i.e. EMGrms, co-activation and EMG mean frequency) confirm that in comparison with isometric contraction alone, isometric contraction with superimposed vibration stimulation induces higher neuromuscular activity in the lower limb muscles. Further, the results also imply strongly and confirm that frequency or amplitude alone does not decide the level of induced neuromuscular activity, and instead the combination of frequency and amplitude along with the level of muscle contraction/tension should be used to identify the ‘optimal’ response to vibratory stimulation.

Both the 30 and 50Hz frequencies were found to elicit significantly higher neuromuscular activity compared to the control in the VL and BF muscles. However, among the vibration variables tested, based on the percentage increases in mean EMGrms activities of the VL and BF muscles, increases in the co-activation (BF/VL and BF/VM) ratios and increases in the MEF values, the 50Hz-0.5mm frequency-amplitude combination was found to be the most effective in generating the highest neuromuscular activity in leg extensors muscles, similarly to previous work on vibrating platforms [39]. This is of particular importance considering that, although previous studies have suggested both 30Hz and 50Hz as suitable stimuli, 50Hz frequency has been shown to be more effective stimulus in lower limbs compared to 30Hz [40]. Also, with regards to the muscle tuning theory discussed earlier, in lower limbs, the highest levels of muscle activity have been observed to coincide with the highest vibration damping which occurred at the resonant frequencies (10-50Hz) of the lower limb tissues [41].

Further, it has also been suggested that the higher frequencies and amplitudes of vibration would be more effective in inducing higher neuromuscular stimulation [42]. However, the results of this study do not indicate that simply delivering a combination of higher frequency and amplitude necessarily induces higher neuromuscular response. The combination of the highest frequency and amplitude stimulation tested during this study (i.e. 50Hz-1.5mm), did not lead to the highest neuromuscular activity compared to other combinations.

A limited number of studies on indirect (WBV, ULV) vibration have compared different combinations of frequency-amplitude stimulation simultaneously for their effectiveness in generating higher neuromuscular activity or muscle strength [42], [43]. To the best of the author’s knowledge, no study has investigated the effect of graded isometric contractions superimposed on vibration in the lower limbs. However, in direct vibration studies, strong evidence specifying the factors that influence neuromuscular response does exist. Increase in muscle length has been linked to increase in TVR [20]. Also, vibration frequency, amplitude and muscle pre-stretch have been specified to influence the TVR [21]. Higher amplitude vibration has led to higher TVR response in animals [44], [45]; potentially due to increased number of muscle-spindle endings being activated leading to increased number of motor neurons being employed [46]. Importantly, previous work has suggested that the higher amplitudes may only be effective in sub-maximal contractions [47]. However, from the results of this study no clear trends indicating higher amplitudes (i.e. 1.5mm compared to 0.5mm) leading to higher neuromuscular responses only under sub-maximal contractions (i.e. 20% to 40% of MVC compared to 80 to 100% of MVC) were found. Sub-maximal contractions did, however, lead to the higher neuromuscular responses with both the lower (0.5mm) and higher (1.5mm) amplitudes in the (VL) muscle. The antagonist muscle (BF) displayed a different response to the VL with a higher neuromuscular activity for MVCs irrespective of the amplitude levels. Without further evidence, it is difficult to infer whether the almost contrasting response of agonist and antagonist is a part of a wider neuromuscular strategy to counteract the vibration perturbation depending on the force level superimposed.

It is also worth noting that the magnitude of acceleration produced by the 50Hz-0.5mm stimulation is equivalent to 30Hz-1.5mm stimulation. Despite having the same acceleration magnitude, the results indicated significant differences between the neuromuscular responses to these vibration stimulations. Further, neuromuscular responses to the same vibration frequency (e.g. 30Hz) differed significantly with the change in the amplitude from 0.5mm to 1.5mm. Overall, the observed differences in neuromuscular responses in this study can be attributed to the combinations of vibration frequencies (30Hz vs. 50Hz), amplitudes (0.5mm vs. 1.5mm) and contraction levels (20 to 100% of MVC). This further affirms the role of the vibration frequency-amplitude combination in grading the neuromuscular response as opposed to the level of acceleration or frequency alone. Nevertheless, from previous evidence and the results of this study it is clear that the role and the effect of vibration amplitude in grading neuromuscular response should not be ignored.

The VM EMGrms response under all the vibration conditions was similar to its control conditions response. This is likely due to the lesser engagement of this muscle in the task used in this study. The VM is likely to be more engaged as an agonist when the knee angle is greater i.e. the knee is more extended. As the length of the muscle and pre-stretch during vibration appear to have direct influence on the neuromuscular response, the 90°knee angle posture employed in this study potentially restricted the involvement of the VM as an agonist, limiting the effect of vibration exercise on the VM’s neuromuscular activity. It is important to note that in the knee extension, the VM acts as a synergist with the VL. In this regard, a recent study suggests that quadriceps muscle activity during leg press exercise depends upon and strongly varies with the knee angle, foot placement and effort level [48]. The VM has been shown also to display a nonlinear EMG/force relationship during isometric leg press exercise [49]. Further, recent investigation which looked at the ratio of VL/VM contraction during knee extension, concluded that the neural drive may be biased towards the VL compared to the VM and seems to be dependent on the force level [50]. The authors suggested that the higher the force generation capacity of an individual’s VL, the higher the bias of neural drive towards VL over the VM. The study also found that this bias increased with increase in the force level during isometric knee extension contraction. The above reasons might explain why, compared to the VL and BF, the VM did not show an increase in neuromuscular activity when stimulated by vibration superimposed on varying levels of contractions.

Contrary to the previous evidence for the upper limb [26], vibration stimulation superimposed on lower contraction levels of 20 to 60% of MVC in this study was found to be equally or more effective in inducing higher neuromuscular activity compared to near maximal/maximal effort levels of 80 and 100% of MVC. In the upper limb study [26], irrespective of muscle group (i.e. agonist or antagonist), higher force levels of 80 and 100% of MVC were able to induce significantly higher EMGrms amplitudes compared to control conditions. In [26] the maximum increase in average EMGrms value was found to be of 77.2%, whereas in the current study, the maximum increase in average EMGrms value was found to be of 118% when compared to the control. Based on the average increases in EMGrms values, vibration superimposed on isometric contraction seems to induce higher neuromuscular activity in the lower limbs compared to that which was reported previously for the upper limbs [26]. This implies upper and lower limb muscles may respond differently to counteract the vibration, possibly with different neural strategies, thus leading to different neuromuscular responses even when stimulated by the same vibration parameters and level of muscle tension.

Although higher neuromuscular response was observed at lower contraction levels for the VL, similar conclusions cannot be drawn about the BF. In fact, the BF showed more significant activity at the higher contraction levels of 80 and 100% of MVC. Compared to the control and with increase in the force level, the VL and BF showed contrasting responses (i.e. the VL converging with the control and the BF diverging with the control). These contrasting responses of the VL and BF could be a neuromuscular strategy to counteract increasing muscle tension when superimposed on vibration. Reasons behind these seemingly contrasting differences in the neuromuscular activity of the upper and lower limb need to be investigated further.

### Co-activation of Agonist and Antagonist

Co-activation of agonist and antagonist muscles at the joint is employed for stabilizing the joint [51]. Indirect vibration stimulation (WBV, ULV) induces a perturbation at the joint [26], [52]. Therefore, when vibration stimulation is delivered, it would be reasonable to expect higher co-activation of the agonist-antagonist pair in order to stabilise the joint. This was indeed the case when co-activation of VL and BF under the vibration condition was compared to the control. In-fact with vibration stimulation, VL-BF showed higher co-activation at almost all the vibration conditions and effort levels.

Under both control and vibration conditions, co-activation levels were higher at lower effort levels and were lowest at the MVC. Interestingly, similar results have been reported in a study conducted on upper limb vibration [26]. The authors of this study [26] argued that when the agonist is involved in lower force production, the joint rotation is primarily controlled by the antagonist hence leading to higher co-activation. The results of our study, also indicate that co-activation of the antagonist may be primarily used as a joint stabilisation mechanism rather than to modulate agonist force output. In this regard, significantly higher co-activation levels (than control) under vibration conditions, at higher force levels may seem contradictory. However, it can be argued that when vibration is superimposed with graded force levels, the higher the force level, the higher the perturbation induced at the joints. Therefore although overall co-activation levels dip at higher force levels, co-activation levels under vibration conditions (at higher force levels of 80 to 100% of MVC) were significantly higher than the respective control conditions.

Under direct vibration stimulation, higher co-activation levels compared to control have been reported [51]. However, extrapolation of the results obtained from direct vibration stimulations to the indirect vibration stimulations should be approached with a caution. Notwithstanding these differences however, co-activation results obtained in this study corroborate earlier findings [51]. In that, vibration stimulation does induce higher co-activation in Agonist-Antagonist pair.

### Potential Mechanisms leading to Increased Neuromuscular Activity

The observed increases in neuromuscular activity under vibration conditions superimposed with graded contraction levels can be ascribed to a range of mechanisms from the local (muscular) to the central (CNS) level.

On the local level, it has been reported that muscles actually damp externally applied vibrations and that activated muscles are capable of absorbing more vibration energy than the muscles in rigor [53], [54]. As a consequence it has been suggested that muscles are activated to attenuate the vibrations [23]. The higher neuromuscular activation levels observed in this study imply that the soft tissue activations to damp the oscillation could have contributed to the observed increases in neuromuscular activity.

The increase in the EMG amplitude also signifies modulation in the motor unit recruitment and/or motor unit discharge frequency. An increase in mean frequency (MEF) can signify the additional recruitment of superficially located high threshold motor units, as these motor units typically contribute large and sharp spikes which influence high frequency bands of sEMG [35], [55]. The enhancement of the contribution to stretch reflexes with indirect vibration stimulation has been attributed to the possible recruitment of high threshold units and muscle fibres [56]. This suggests that vibration could potentially modify motor unit recruitment patterns and rate coding behaviour, possibly recruiting high threshold motor units leading to enhancement in neuromuscular responses.

In addition, it is known that direct vibration stimulation induces TVR by stimulating primary and secondary afferents [1], [44]. Discharge of these afferents have been reported to be dependent on muscle pre-stretch, and the discharge increases with increase in muscle stretch-length [1]. Whereas voluntary isometric contraction also increases this discharge [1]. Vibration also stimulates, Ib afferents from Golgi Tendon Organ [1], [57] and Ib afferents are stimulated more when muscle contracts [44]. Thus, vibration has the ability to alter significantly the sensitivity of primary, secondary afferents and Ib afferents leading to an increase in neuromuscular response. Muscle length and isometric contraction seem to have direct effect on the spindle sensitivity altering the neuromuscular response further. Considering our observations, it could be that the increased neuromuscular activity observed with the superimposition of vibration may have been results of alterations in afferent responses due to vibration.

### Limitations of the Study

As discussed earlier, it is still not clear whether the electromyography amplitude response found to be synchronous with vibration stimulation frequency is due to motion artifacts or is the result of stretch reflex response [18], [36]. Due to this uncertainty, no artifact removal processing (except at 50Hz) was performed on the EMG data obtained in this study. This might be considered a limitation. However, 50Hz line interference was observed in all of the WBV EMG data and a notch filter centred at 50Hz frequency was used to attenuate the line interference during both 30Hz and 50Hz stimuli EMG data. It is important to note that of all the frequency amplitude stimuli combinations tested for this study, 50Hz stimulation with (0.5mm amplitude) induced the largest neuromuscular activity. And overall, 50Hz stimulations induced equal or higher neuromuscular activity compared to 30Hz stimulations.

If it is assumed that vibration stimulation leads to motion artifacts in the sEMG data at the frequency of vibration stimulation and harmonics thereof, the largest energy of the so called ‘motionartifact’ is concentrated at the stimulus frequency [36]. For the EMG data collected for this study, the spike at 50Hz were amongst the largest (although it did not necessarily contain high energy). Despite removing the most significant proportion of the possible ‘motion artifact’ (with the largest spike) around 50Hz, from the EMG data, the 50Hz stimulus led to equal or higher neuromuscular activity compared to the 30Hz in this study. Further, despite the fact that, the signal at 30Hz frequency was not removed from the 30Hz stimulation sEMG data, the general trends of 30Hz and 50Hz neuromuscular responses (i.e. EMGrms, MEF) were quite alike. This gives further confidence in the results of this study, in that, the possibility of motion artifact skewing the EMG data and the results is quite limited.

It is important to note that in the vibration exercise superimposed with graded isometric contraction, transmission of the vibration through the limbs would be dependent on muscle contraction [16], [58]. Thus, the degree of muscle contraction and body posture (e.g. knee angle) would in effect dictate the level of vibration transmission and this could have implications on EMG artifacts. Thus motion artifact/stretch reflex responses might be different when WBV is combined with graded isometric contractions. Hence to analyse the motion artefact or stretch reflex’s presence, dedicated and specific signal processing methods may need to be devised [59] and adapted according to the variables (e.g. force/contraction level and stimuli characteristics etc.) specific to the study.

## Conclusions

(A) Isometric contraction superimposed on vibration stimulation leads to higher neuromuscular activity compared to isometric contraction alone in the lower limbs.

(B) In the agonist muscles during a leg press task, vibration exercise with lower contraction levels of 20 to 60% of MVC force seem to generate higher neuromuscular activation compared to the higher levels of 80 to 100% of MVC.

(C) In the antagonist, higher contraction levels of 80 to 100% seem to induce equal or more neuromuscular activity compared to the lower contraction levels. Whether this apparently contrasting difference between the agonist and the antagonist responses at higher contraction levels of 80 to 100% of MVC is a part of wider neuromuscular response strategy is unclear.

(D) Among the vibration variables tested, the 50Hz-0.5mm stimulus generated the highest neuromuscular response compared to the control irrespective of the muscle group and/or contraction level.

(E) Both 50Hz and 30Hz, frequencies led to higher neuromuscular activity compared to the control however, the combination of the frequency with the amplitude and the muscle tension together seem to grade the final neuromuscular output instead of frequency alone.

(F) Compared to the control, vibration stimulus led to higher agonist-antagonist co-activation in all conditions and effort levels except 20% of MVC.

(G) Sub-maximal and maximal levels of 80 and 100% MVC contraction force led to the most significant co-activation difference between the control and the vibration

## Authors’ contributions

MC, RDN and ANP conceptualised and designed the study. MC and RDN acquired the funding. RDN developed the initial design of the device. ANP and RDN further developed the device. ANP completed the hardware, instrumentation and computer interfacing with the software programming required for the device operation. ANP acquired the data and processed and analysed the data from the study. ANP interpreted results from the data and RDN and MC provided critical input to the results. ANP drafted the original manuscript. ANP finalized the manuscript. All authors read and revised the manuscript critically for important intellectual content, and approved the final manuscript for publication. All authors agree to be accountable for all aspects of the work.

## Acknowledgements

Authors would like to thank Scottish Funding Council (SFC) for funding to support the study. The authors thank the volunteers for their participation. ANP would also like to thank Royal Society of Edinburgh for the J M Lessells Travelling Fellowship and to Prof. Kevin Englehart of University of New Brunswick, Canada for his expert advice during the fellowship in relation to this work.

## Availability of data and materials

Data and materials can be made available upon request to the authors.

## Ethics approval and consent to participate

Experiments were approved by the North of Scotland Research Ethics Committee (NOSREC) of the National Health Service (NHS) Grampian, Aberdeen, UK.

## Consent for publication

Not applicable

